# The ectromelia virus virulence factor C15 facilitates early viral spread by inhibiting NK cell-infected cell contacts

**DOI:** 10.1101/2022.06.24.497501

**Authors:** Elise M. Peauroi, Stephen D. Carro, Luxin Pei, Glennys V. Reynoso, Heather D. Hickman, Laurence C. Eisenlohr

## Abstract

The success of poxviruses as pathogens depends upon their antagonism of host responses by multiple immunomodulatory proteins. The largest of these expressed by ectromelia virus (the agent of mousepox) is C15, one member of a well-conserved poxviral family previously shown to inhibit T cell activation. Here, we demonstrate by quantitative immunofluorescence imaging that C15 also limits contact between natural killer (NK) cells and infected cells *in vivo*. This corresponds to an inhibition in the number of total and degranulating NK cells, *ex vivo* and *in vitro*, with no detectable impact on NK cell cytokine production nor the transcription of factors related to NK cell recruitment or activation. Thus, in addition to its previously identified capacity to antagonize CD4 T cell activation, C15 inhibits NK cell cytolytic function, which results in increased viral replication and dissemination *in vivo*. This work builds on a body of literature demonstrating the importance of early restriction of virus within the draining lymph node.

**summary:** Poxvirus B22 family proteins are important virulence factors known to inhibit T cell functions. Peauroi et al. identify a novel function of the ectromelia virus homolog, C15, which inhibits NK cell-target contact and cytolytic function to facilitate early viral spread.

(Provide a short, ∼40-word summary statement for the online JEM table of contents and alerts. This summary should describe the context and significance of the findings for a general readership; it should be written in the present tense and refer to the work in the third person.)

## Introduction

The extraordinary virulence of poxviruses can be attributed to their extensive antagonism of the host immune response (reviewed in: Smith and Kotwal, 2002; Sigal, 2016). The orthopoxvirus (OPXV) genus includes variola virus (VARV, the agent of smallpox), vaccinia virus (VACV, the gold standard smallpox vaccine) and multiple viruses that pose threats from zoonotic transmission including monkeypox virus (MPXV). OPXVs are the largest mammalian viruses with genomes containing around 200 open reading frames (ORFs) (Buller and Palumbo, 1991). Many of these ORFs encode immunomodulatory proteins that are exquisitely tuned to the immune response of the host. Thus, their functions are best studied during natural host-pathogen relationships. Mousepox, the murine poxviral disease caused by ectromelia virus (ECTV), provides a unique opportunity to study natural OPXV infection in a tractable system (Chapman et al., 2010).

One notable ECTV immunomodulatory protein is C15. It is a member of the B22 protein family, which are the largest OPXV proteins at nearly 2000 amino acids. B22 family members are well conserved across the virulent OPXVs (including VARV, MPXV and ECTV), but the C15 ORF is truncated in VACV. The size and conservation of B22 family members suggest a large contribution to virulence, and this has been demonstrated for the homologs in MPXV and ECTV (Alzhanova et al., 2014; Reynolds et al., 2017; Forsyth et al., 2020). Functionally, B22 family members have been shown to modulate T cell responses via complex mechanisms. The VARV, MPXV and cowpox virus (CPXV) B22 homologs inhibit primate CD4 and CD8 T cells, but B22 family proteins belonging to OPXV that infect murine cells (CPXV and ECTV) selectively inhibit CD4 but not CD8 T cell function (Alzhanova et al., 2014; Forsyth et al., 2020). These data suggest that primate and murine CD8 T cells differ in ways that dictate susceptibility to B22-mediated inhibition. While the specific mechanism of this inhibition has not been revealed, our previous work with ECTV demonstrated that C15 interferes with the formation of CD4 T cell synapses (Forsyth et al., 2020).

C15 inhibitory function appears to extend beyond adaptive immunity. Work by others indicated that C15 may impact viral spread in the draining lymph node as early as 4 days post infection (dpi) of susceptible BALB/c mice (Reynolds et al., 2017), suggesting antagonism of the innate immune response that contributes to virulence. Natural ECTV infection of mice is modeled by footpad injection, and the first stages of the host response in the draining popliteal lymph node (LN) have been well characterized in C57BL/6 (B6) mice, which are resistant to lethal ECTV infection (Fang et al., 2008; Xu et al., 2015; Wong et al., 2018, 2019; Ferez et al., 2021; Melo-Silva et al., 2021). Natural killer (NK) cells are critical mediators of early protection and are necessary for resistance to ECTV infection in the first 4 dpi (Fang et al., 2008). NK cells contribute to viral control through the production of antiviral IFNγ and by perforin-mediated cytolysis (Parker et al., 2007; Fang et al., 2008). Previous work has revealed the mechanisms of NK cell recruitment to the infected LN (Xu et al., 2015; Wong et al., 2018, 2019), of early NK cell activation to produce IFNγ (Wong et al., 2018) and of NK cell recognition of infected cells leading to cytolysis (Fang et al., 2008, 2011; Ferez et al., 2021). Due to their importance in the early control of infection, NK cell function is targeted by many viruses, including OPXV, to promote viral spread (reviewed in: Burshtyn, 2013; Ma et al., 2016, 2021). Examples include the OPXV MHC class I-like protein (Campbell et al., 2007) and hemagglutinin protein (Jarahian et al., 2011), which interfere with NK cytolysis by inhibiting NK cell activation, and multiple mechanisms of antagonism of the NK cell-activating cytokine IL-18 by ECTV (Esteban and Buller, 2004; Melo-Silva et al., 2011).

Few viral proteins have been reported to antagonize the function of both innate (NK cell) and adaptive (T and B cell) lymphocytes (for example: Krmpotić et al., 2002). Here, we demonstrate that the B22 family member of ECTV, C15, inhibits NK cell cytolytic function by limiting target cell contact in addition to its previously identified capacity to antagonize CD4 T cell activation. The impact of this inhibition allows for increased viral replication and dissemination within the draining LN early after infection.

## Results

### C15 facilitates early ECTV replication and dissemination in an NK cell-dependent manner

To investigate the apparent impact of C15 on early stages of ECTV infection (Reynolds et al., 2017), we analyzed viral titers in the draining popliteal LN at 1-3d after infection of B6 mice with eGFP-expressing WT ECTV (WT) and C15-deficient ECTV (ΔC15). Expression of C15 profoundly increased viral titers in the LN by 3 dpi, evident both by traditional titering (Fig. S1 A) and focus forming assay (Fig. 1 A). Given the magnitude of the impact at these early timepoints, we hypothesized that C15 targets an aspect of innate immune control.

**Figure 1:**
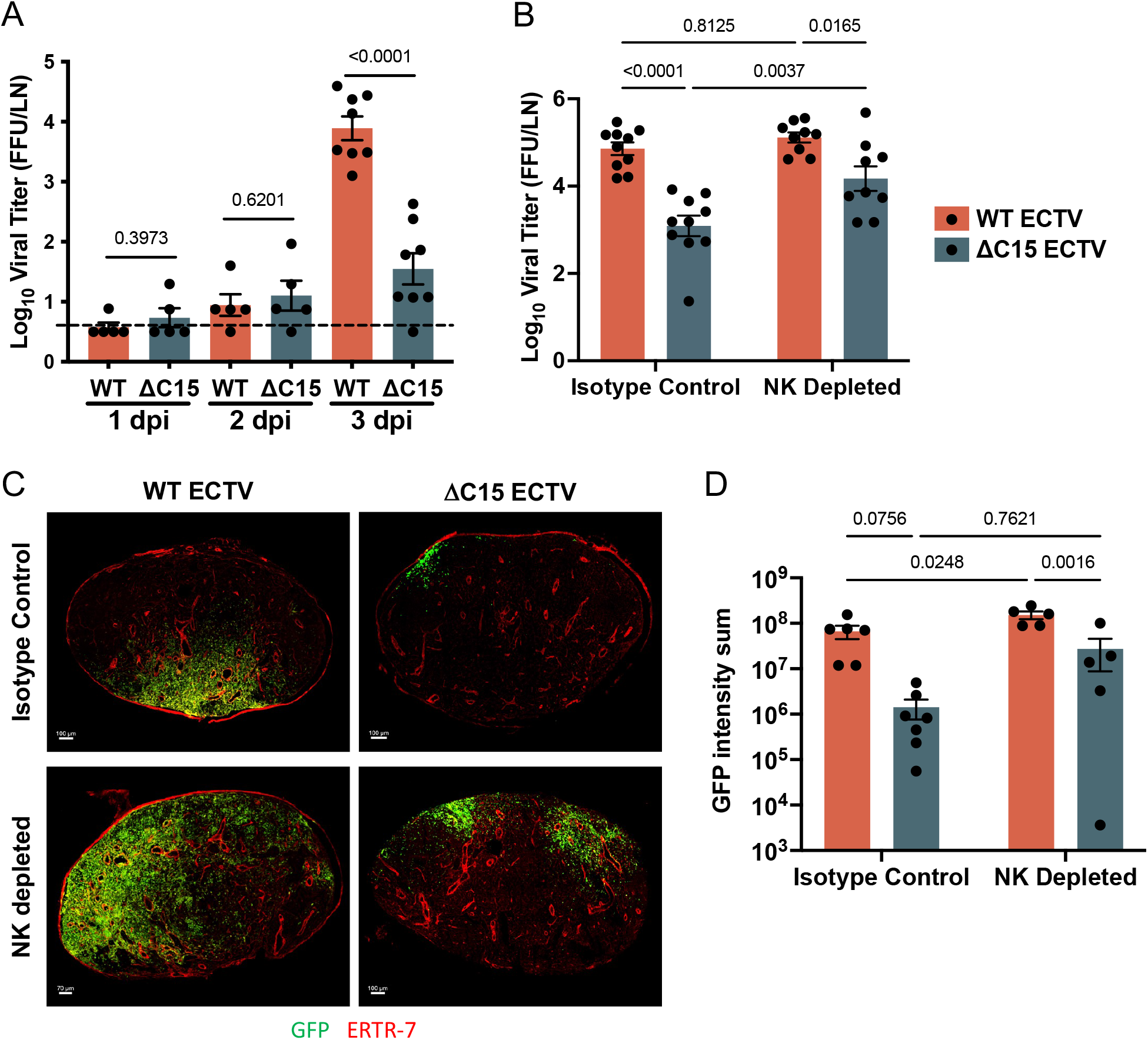
C15 facilitates early ECTV replication and dissemination in an NK cell-dependent manner. **(A)** C57Bl/6 (B6) mice (days 1 and 2: n=5, single experiment. Day 3: n=9 pooled from 3 separate experiments titered together) were infected with 3000 pfu of WT or ΔC15 ECTV, sacrificed 1-3 dpi and popliteal LNs were harvested. **(B-D)** B6 mice were injected i.p. with either anti-NK1.1 (NK depleted) or an isotype control antibody and the next day were infected by footpad injection with 3000 pfu of eGFP expressing WT or ΔC15 ECTV, sacrificed 3 dpi and popliteal LNs were harvested. The LNs were analyzed as follows: **(A and B)** LNs were homogenized and viral titers were determined by focus forming assay on BSC-1 cells. Graphs display log_10_ transformed titers as focus forming units (FFU)/LN. Bars = mean, Error bars = SEM. T Tests (A) or a 2-way ANOVA with multiple comparisons was performed (B), p values shown. (B: n=9-10 per group pooled from 3 separate experiments titered together). **(C and D)** LNs were sectioned and analyzed for the indicated markers by immunofluorescence analysis, virus infected cells are identified by GFP staining and ERTR7 (a fibroblastic reticular cell marker) demonstrates the LN stroma. (n=5-7 from 3 separate experiments). **(C)** Representative images, maximum intensity projections with 70-100 μm scale bars (indicated on image). **(D)** Level of virus infection in each section was determined by the sum of the GFP signal, shown on a log_10_ scale. Bars = mean, Error bars = SEM. A 2-way ANOVA with multiple comparisons was performed, p values shown.

NK cells provide critical early control of ECTV infection (Parker et al., 2007; Fang et al., 2008); therefore, we next asked whether C15 targets this cell type to gain a replicative advantage. We depleted B6 mice of NK cells using anti-NK1.1 antibody (Fig. S1 B) and found that, in comparison to isotype control-treated mice, NK cell depletion largely eliminated the replicative advantage conferred by C15 expression (Fig. 1 B). Additionally, NK depletion did not statistically impact viral titers in the LN of mice infected with WT virus, whereas replication of ΔC15 virus was significantly increased in the absence of NK cells (Fig. 1 B). These results suggest that C15 acts on NK cells to facilitate replication at this early timepoint.

We were next interested in identifying whether C15 impacts viral dissemination within draining LNs of B6 mice. We harvested LNs 3 dpi with WT or ΔC15 ECTV, imaged these tissues by confocal microscopy and analyzed the amount and distribution of infected cells (GFP+) in relation to the LN stroma (ERTR7+) (Figs. 1, C and D). Mirroring the titer data, there was an increase in GFP expression in NK-intact (isotype control-treated) mice infected with WT vs ΔC15 (Figs. 1, C and D), and we noted differential distribution of GFP+ virus-infected cells in the two infection conditions (Fig. 1 C). Similar to previous observations in BALB/c mice (Reynolds 2017), the viral GFP signal in NK-intact B6 mice was restricted to the LN periphery in ΔC15 infection compared to a larger area of GFP expression during WT infection, suggesting that C15 facilitates increased viral replication and spread *in vivo*.

We also performed this analysis in NK cell-depleted mice and found that, mirroring the titer data in Fig. 1 B, the difference in GFP expression between WT and ΔC15 infection was lessened in NK-vs isotype control-depleted animals (Fig. 1 D), further supporting the conclusion that C15 exerts an immunomodulatory function on NK cells. In NK cell-depleted mice, visual analyses showed that depletion provided a more disseminated phenotype even in the absence of C15 expression (Fig. 1 B), though infection disrupted the LN architecture precluding categorization of nodal distribution. Considering the variation between biological replicates, there appeared to be increased nodal spread in WT infected tissues, both NK cell-depleted and -intact, compared to NK cell-intact ΔC15 infected LNs (Figs. 1 C, S2), suggesting that NK cells are able to restrict virus specifically in the absence of C15 expression. These data are consistent with targeting of NK cell responses by C15 to facilitate early viral replication and dissemination in the LN and suggest that C15 is a major contributor to ECTV-mediated inhibition of NK cells. Our findings are consistent with previous work (Reynolds et al., 2017) showing that during ΔC15 infection, increasing the inoculating dose decreased titers in all central sites but not at the site of infection (contrary to WT infection where dose was correlated with titer at all sites), suggesting that in ΔC15 infection, increased viral loads are more easily detected by the host and spread is better controlled by the immune response.

### Viral expression of C15 reduces numbers of total and degranulating NK cells in infected tissue

In order to assess the impact of C15 on the NK cell response *in vivo*, we next analyzed WT-vs ΔC15-infected LNs 3 dpi by flow cytometry. Despite the substantial increase in virus present in WT vs. ΔC15 infected tissues at this time (Fig. 1 A), there was no difference in the total number of live cells recovered from the two infection conditions (Fig. 2 A). Nevertheless, NK cells made up a significantly increased percentage of the live cells in the LN during ΔC15 infection (Fig. 2 B) and there was a similar, though less profound, difference in the absolute number of NK cells identified (Fig. 2 C). We also analyzed the percentage and number of an unrelated innate cell type, F4/80^+^ macrophages, which followed the opposite trend, where there was a similar number (Fig. 2 C) but increased percentage (Fig. 2 B) in WT vs ΔC15 ECTV infection; this trend fits more closely with the expected immune cell response based on viral titer alone.

**Figure 2:**
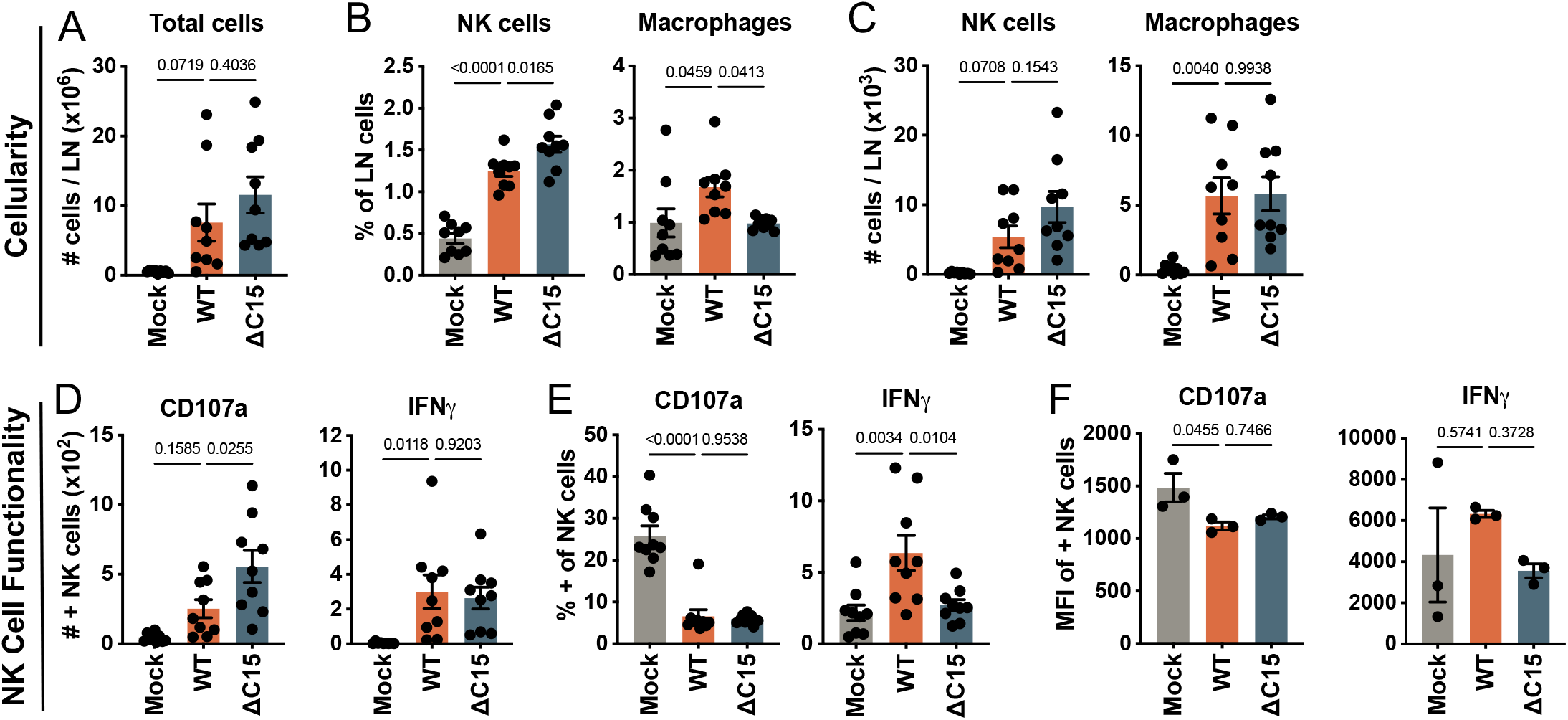
Viral expression of C15 reduces numbers of total and degranulating NK cells in infected tissue. B6 mice were infected by footpad injection with 3000 PFU of GFP-containing WT or ΔC15 ECTV or mock infected with PBS and sacrificed 3 dpi. The draining popliteal LNs were harvested, incubated in monensin for 1 hour, then dissociated and cells were stained and analyzed by flow cytometry. **(A)** Total number of cells isolated from each LN. NK cells were gated as NK1.1+CD3-and **(B)** total number and **(C)** percent of this population and of F4/80+ macrophages are shown. The number **(D)**, percent **(E)** and median fluorescence intensity (**F**, MFI, single representative independent experiment shown) of NK cells expressing CD107a on the surface and IFNγ intracellularly. n=9 per group, data pooled from 3 independent experiments. A one-way ANOVA with multiple comparisons was performed and p values are reported. Bars = mean, error bars = SEM.

We next assessed the functional responses of NK cells in the LNs of WT-and ΔC15-infected mice by flow cytometry. Surface CD107a staining is a marker for cytolytic degranulation; there was a significant increase in the number of CD107a+ NK cells in ΔC15 vs WT infection (Fig. 2 D), but notably, there was no change in the proportion of CD107a+ degranulating NK cells between infection conditions (Fig. 2 E). This suggests that the difference in NK cell cytolytic activity is directly attributable to the increased frequency of NK cells in the LN. There was also no difference in the median fluorescence intensity (MFI) of CD107a on positive cells (Fig. 2F), suggesting that C15 does not impact NK cell degranulation on a per cell basis.

NK cells not only act as important cytolytic effectors, but are also crucial early sources of IFNγ, which orchestrates the antiviral immune response to ECTV in the LN (Fang et al., 2008; Xu et al., 2015; Wong et al., 2018). Although there was no difference in the number (Fig. 2 D) or the MFI (Fig. 2 F) of IFNγ+ NK cells in WT vs ΔC15 infection, the percentage of IFNγ+ NK cells was higher in WT vs ΔC15 infection (Fig. 2 E), in marked contrast with CD107a. This is consistent with the finding that early IFNγ production by NK cells is in response to infected cells which are increased in WT vs ΔC15 infection (Wong et al., 2018). Thus, C15 does not impact NK cell production of IFNγ or degranulation on a per cell basis; instead, C15 reduces the total number of NK cells and thereby the number of degranulating NK cells in infected LNs.

### C15 does not impair transcription of factors involved in NK cell recruitment and activation

We next sought a potential explanation for the increased number of NK cells in

ΔC15 infected tissues 3 dpi by assessing earlier transcriptional differences in molecules involved in NK cell activation and recruitment. We collected total RNA from infected LNs 40 hours pi, a time before obvious divergence of virus replication by titer (Fig. 1 A) and when we identified the first infected cells in the LN by imaging. Reverse transcription followed by quantitative PCR revealed significant increases in virus, indicated by the viral transcript *Evm003*, in WT vs ΔC15 infected LNs even at this early time (Fig. 3 A). This demonstrates that C15 facilitates viral replication at very early times post infection.

**Figure 3:**
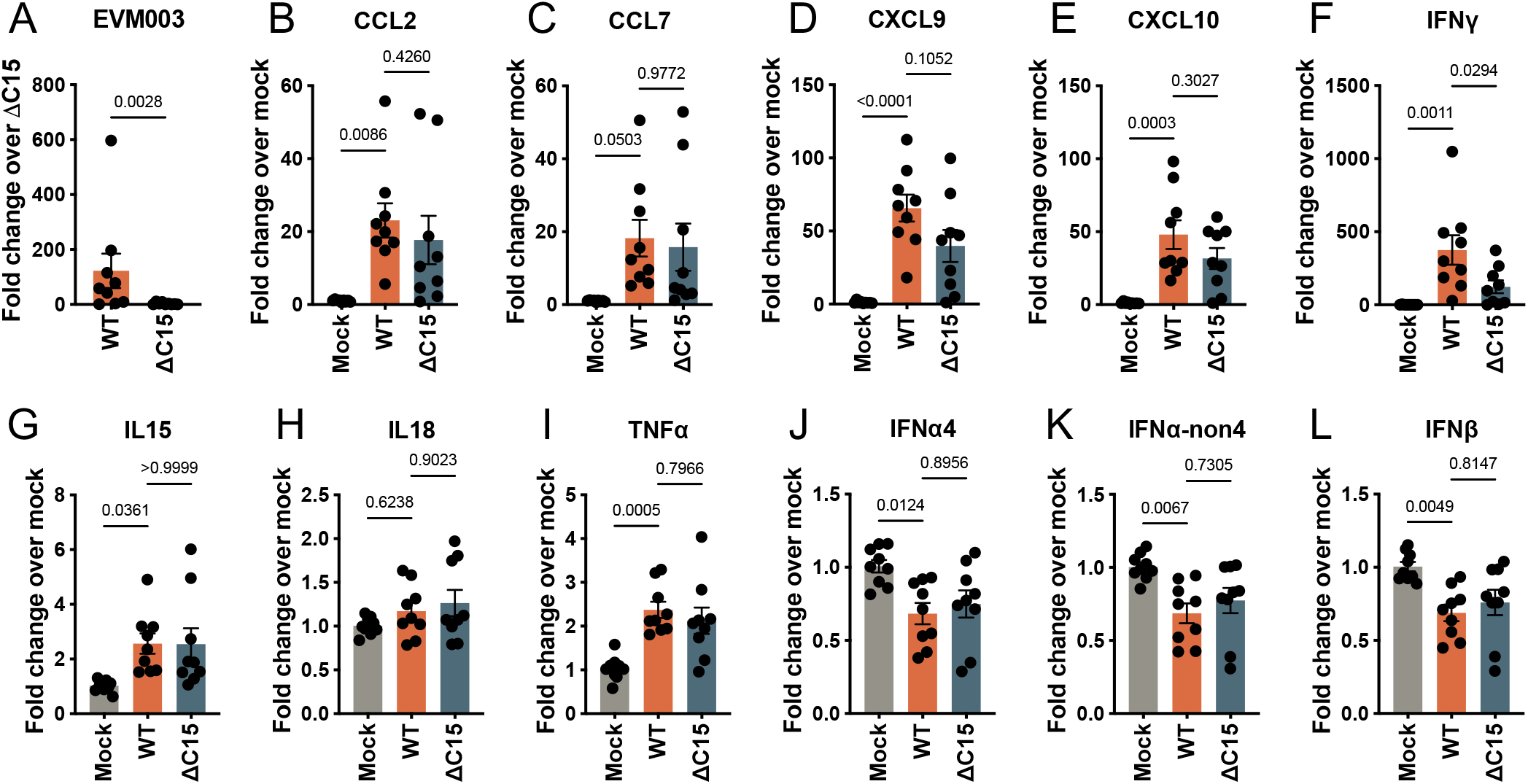
C15 does not significantly inhibit transcripts for cytokines, chemokines or interferons on a whole tissue level. Popliteal LNs were harvested 40 hours post infection of B6 mice infected with WT ECTV, ΔC15 ECTV or PBS (mock) and expression of mRNA for viral *Evm003* **(A)**, the chemokines *Ccl2* **(B)**, *Ccl7* **(C)**, *Cxcl9* **(D)**, *Cxcl10* **(E)**, the inflammatory cytokines *Ifng* **(F)**, *Il15* **(G)**, *Il18* **(H)**, *Tnfa* **(I)**, *Ifna4* **(J)**, *Ifna-non4* **(K)** *and Ifnb* **(L)** are analyzed relative to host gene *Gapdh* expressed as fold change. *Il12* and *Perforin* were also analyzed but were often below the limit of detection and therefore data is not shown. Measured by RT-PCR, points are biological replicates (n=9) calculated as the average of two technical replicates and pooled from three independent experiments, each normalized to the mean of the ΔC15 ECTV-(A) or mock-(B-L) infected samples. A Mann-Whitney test (A) or one-way ANOVA with multiple comparisons (B-L) was performed; p values are shown. Bars = mean, error bars = SEM

We then analyzed the transcription of various cytokines and chemokines reported to be important during the early stages of ECTV infection and relevant to NK cell recruitment to ECTV-infected tissue and to NK cell activation. Recruitment of NK cells to the LN involves a coordinated response by different cell types beginning with infection-induced CCL2 and CCL7 production leading to increased IFNγ and finally CXCL9-and CXCL10-mediated recruitment of circulating NK cells to the LN (Xu et al., 2015; Wong et al., 2018, 2019). While chemokine transcription was clearly induced by infection, we detected no statistically significant increases in transcription of selected chemokines in the absence of C15 (Figs. 3, B-E). Consistent with our findings by flow cytometry (Figs. 2 D-F) and aligning with the increase in detected viral transcripts (Fig. 3 A), we detected a significant increase in IFNγ transcripts in WT infection (Fig. 3 F), again demonstrating that C15 does not inhibit the IFNγ response at this time. C15 had no impact on the transcription of NK cell-activating cytokines IL-15 and IL-18 (Figs. 3, G and H), the NK cell effector molecule TNFα (Fig. 3 I) or the type I-IFNs (Figs. 3, J-L) which are also important NK cell activators (Melo-Silva et al., 2021). Overall, there were no instances in which transcripts increased in ΔC15 vs. WT infection. Therefore, we conclude that C15 does not directly target the transcription of factors in the LN involved in NK cell recruitment or activation. Results of the transcript analyses (Fig. 3) together with the *ex vivo* flow data (Fig. 3) support a model in which C15 selectively protects infected cells from NK cytolysis.

NK cell cytolysis and IFNγ production are differentially regulated. NK cells can be activated by cytokines, which signal through the JAK-STAT pathway leading to IFNγ production and promoting cell replication. Separately, NK cells recognize target cells at contact points called immunological synapses by the integration of NK cell activating and inhibitory receptors through DAP-10/DAP-12 adaptors; this can trigger cytolysis, proliferation and IFNγ release via a mechanism more similar to T and B cell receptor signaling (Brandstadter and Yang, 2011; Piersma and Brizić, 2021). Therefore, while IFNγ production can be triggered by both cytokine-mediated activation and by target recognition, NK cell cytolytic degranulation occurs only following contact with a target cell resulting in formation of an activating immunologic synapse. Since we determined that C15 reduces the number of degranulating NK cells (Fig. 2 D) but does not inhibit the transcripts for NK cell-activating cytokines by RT-PCR (Figs. 3 G-L) or inhibit IFNγ by flow cytometry (Figs. 2 D-F) or RT-PCR (Fig. 3 F), our data support a model in which C15 interferes with the target cell contact-dependent activation of NK cells.

### C15 inhibits NK cell degranulation *in vitro*

In order to test our model, we sought to investigate whether the expression of C15 on infected cells impacts their ability to be killed by NK cells *in vitro*. Due to viral cytopathic effects, inhibition of NK cells by other ECTV immune modulators (Burshtyn, 2013), and poor infectability of traditional murine target cell lines, we were unable to achieve reliable results when using traditional readouts of target cell death. Therefore, we explored differences in the functions of NK cells following co-culture with infected targets *in vitro* by evaluating surface CD107a staining and intracellular IFNγ production.

We used a B6-derived primary skin fibroblast line as targets and found that these cells induced high levels of NK cell degranulation at baseline, but that this was selectively inhibited by C15-expressing WT infected target cells (Fig. 4 A). In contrast, and concordant with our model, we saw no significant differences in the production of IFNγ by NK cells when co-incubated with WT or ΔC15 infected targets (Fig. 4 A). We also performed this assay in the presence of blocking antibody for NKG2D, an NK cell activating receptor that has been shown to be necessary for optimal NK cell responses to ECTV infection (Fang et al., 2008). We found that blocking NKG2D inhibited the overall NK cell response to target cells, consistent with previous findings (Fang et al., 2008), but that it had no impact on the inhibitory effect of C15, suggesting that C15 does not target NKG2D-mediated NK cell activation (Fig. S3 A). Although the effect is slight in comparison to the degree of impact seen *in vivo*, these data demonstrate that expression of C15 on target cells results in a significant inhibition of NK cell cytolytic degranulation and support our model in which C15 selectively interferes with NK cytolysis.

**Figure 4:**
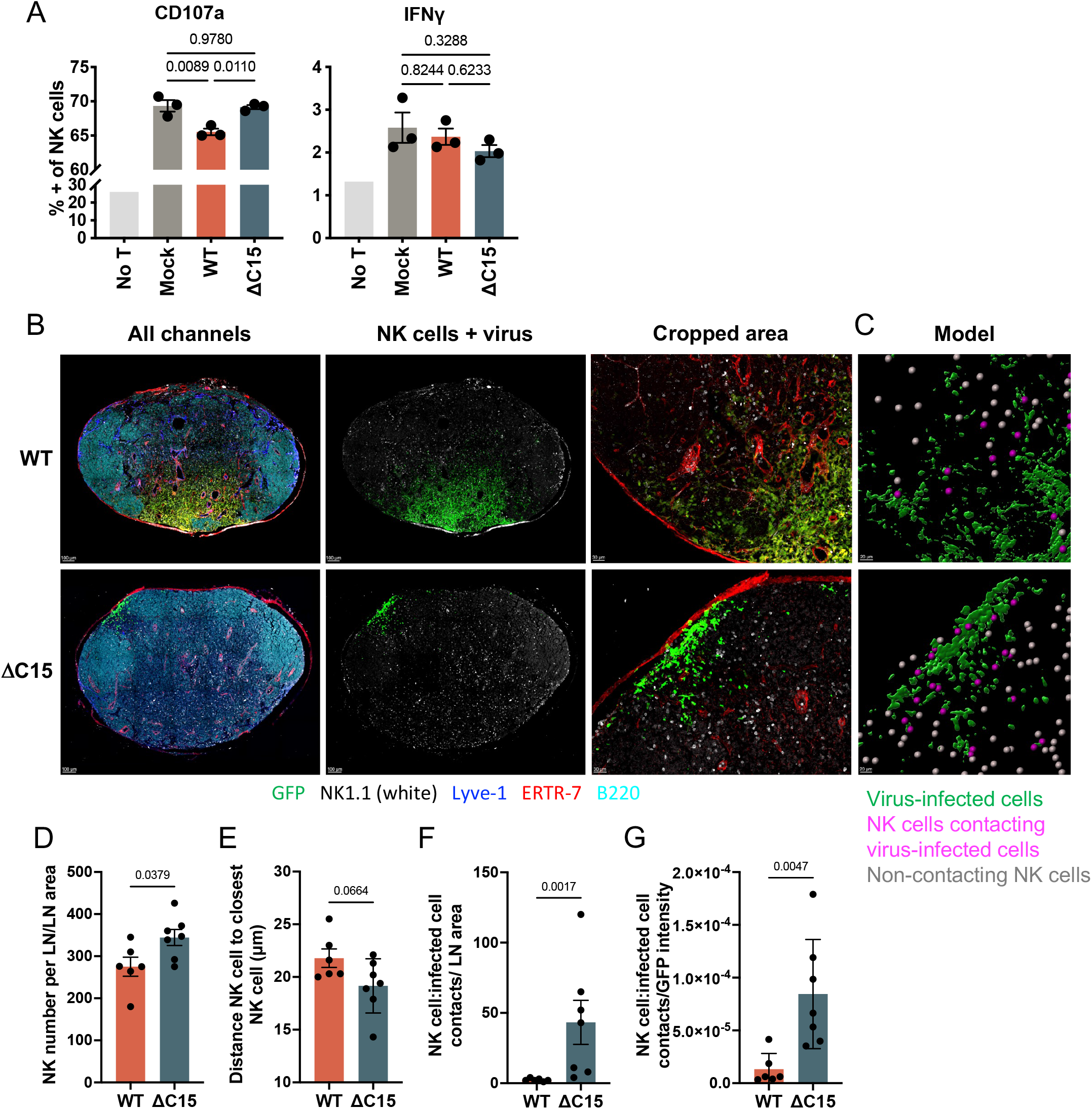
C15 inhibits NK cell degranulation and interaction with virus infected cells. **(A)** Murine fibroblasts were infected with GFP-containing WT or ΔC15 ECTV or mock infected with PBS for 6 hours. They were then coincubated in technical triplicate at a 1:4 effector to target ratio in the presence of monensin and anti-CD107a antibody with NK cells purified from the spleen of a single B6 mouse 6 dpi with 3000 PFU of ΔC15 ECTV, or these NK cells were incubated alone without targets (No T). After 5 hours of coincubation, the cells were stained intracellularly for IFNγ and then analyzed by flow cytometry for the proportion of NK cells displaying the two functions. A one-way ANOVA with multiple comparisons was performed, p values are shown. Data points are technical replicates, bars = means, error bars = SEM. Representative of 4 independent experiments. **(B-D)** The same images from Figs. 1, B and C were analyzed as follows: The following markers discriminate LN anatomic regions: Lyve-1 (a lymphatic endothelial cell marker highlighting lymphoid sinuses), ERTR7 (a fibroblastic reticular cell marker delineating the LN stroma) and B220 (a B cell marker to demonstrate follicles). Virus infected cells are visualized by staining for eGFP and NK cells are identified as NK1.1 surface positive. **(B)** Representative images demonstrating generation of **(C)** a model of NK cells and virus infected cells. This model was used to calculate **(D)** the number of NK cells in each section normalized to LN section area, **(E)** the average distance between each NK cell and the next closest NK cell in μm, and the number of NK cells contacting virus infected cells (threshold = 5 μm, magenta in model) normalized to **(F)** LN section area or **(G)** the viral GFP signal. Bars = mean. Error bars = SEM. T tests were performed, p values shown. Scale bars are labeled, ranging from 15-100 μm.

### C15 inhibits NK cell contacts with virus-infected cells

Given that our data point to a selective impact of C15 on NK cell cytolysis, which is a function dependent upon contact between NK cells and target cells at an immunological synapse, we analyzed NK cell interactions with infected cells by immunofluorescence imaging of LNs. Using the same images as in Figs. 1 and S2, cell-surface NK1.1+ cells were first identified using automated computer imaging analysis software and then classified according to the distance of each individual cell from GFP+ virus-infected cells (Figs. 4, B and C). When normalized to the total LN area, we observed an increase in the overall number of NK cells in ΔC15 vs WT infected LNs, despite the restricted replication of virus (Fig. 4 D), recapitulating our findings by flow cytometry (Fig. 2 B). We then measured the distance between each NK cell and the next closest NK cell, finding that C15 does not substantially impact the overall distribution of NK cells within the LN (Fig. 4 E). To evaluate the interaction of NK cells with infected cells, we next compared the distance of NK cells to sites of infection between conditions, classifying cells located within < 5 μm of each other as contacting (Fig. 4 C). This revealed that C15 profoundly decreased the number of NK cells contacting infected cells when normalized both by LN area and by GFP expression (Figs. 4, F and G). Overall, we conclude that C15 limits NK cell contacts with ECTV-infected cells, decreasing opportunities for activation-induced cytolytic degranulation.

Our imaging analyses highlight the value of spatial data in studies of viral pathogenesis. Due to significant biologic variation during this natural murine infection, we found the data implicating NK cells as a target of C15 were less clear by titer (Fig. 1 B) than when visualized in the draining LN by immunofluorescence imaging (Fig. 1 C). Even when the magnitude of viral GFP expression was low, we still observed an increased distribution of virus in the presence of C15 or in the absence of NK cells. Similarly, imaging provided crucial spatial context in our analysis of the NK cell response. Using imaging, we detected a profound difference in the contacts between NK cells and virus-infected cells (Figs. 4, F-G) in comparison to the more modest difference in the overall number of NK cells identified in the LN by imaging (Fig. 4 D) as well as by flow cytometry (Figs. 2 B-C). In whole organ analyses such as flow cytometry, the decrease in viral replication due to absence of a virulence factor confounds detection of an increase in the response that the factor inhibits. In contrast, in our imaging analysis, the spatial context instead highlighted that inverse relationship and provided potential mechanistic insight, allowing us to observe that C15 profoundly decreases NK cell-target cell engagement proportional to the amount of virus (Fig. 2 G). We also used an *in vitro* assay to confirm that C15 selectively inhibits NK cell degranulation. Based on the difficulty we had using assays that read out target lysis, we resorted to an *in vitro* NK cell degranulation assay but found that there was a high level of background degranulation in response to uninfected target cells and only a modest inhibitory effect of C15 (Fig. 4 A). We conclude that that this assay does not accurately reveal the full-scale impact of C15 on NK cells and speculate that traditional cytolysis assays are not optimized to reveal differences in the context of infected targets. There is a need for *in vitro* assays better able to reveal impacts of virally expressed factors on NK cell cytolysis and accurately predict their effects on NK cell biology *in vivo*.

Overall, we demonstrated that C15 selectively impacts NK cell contact with target cells and cytolytic function without impacting IFNγ production, both *in vitro* and *ex vivo*, and that this facilitates early viral replication and spread *in vivo*. Our data support a model in which C15 protects infected cells by interfering with engagement of target cells and subsequent activating/inhibitory receptor-based activation of NK cell cytolysis. We eliminated NKG2D (an activating receptor important to NK control of ECTV and a well-studied viral target (Ma et al., 2016, 2021)) as the mechanism (Fig. S3 A), but there are many other activating and inhibitory NK cell receptors that C15 could target. We found that WT ECTV had no early replicative advantage in β2 microglobulin-deficient mice (Fig. S3 B), suggesting a nonclassical MHC class I-based mechanism of NK cell activation as a potential target of C15.

The findings shown here are particularly interesting when considered in the context of what else is known about B22 family proteins. With the addition of this novel NK-inhibitory function, we have now revealed that in murine cells C15 is able to inhibit NK and CD4 T cell but not CD8 T cell contact and function. While the specific molecular mechanism of C15 antagonism of CD4 T cells remains to be determined, it appears to impact the formation of immunological synapses (Forsyth et al., 2020). Although there are significant distinctions between T cell and NK cell synapses, these contact-dependent interaction sites are also implicated by results shown here. Together, our data prompt a nuanced exploration of how B22 family proteins selectively impact CD4 and CD8 T cell and NK cell activation. Further work is necessary to explore the molecular mechanism(s) of this protein, which will reveal important underlying biology about similarities and differences of lymphocyte immunological synapses.

## Materials and Methods

### Mice and infections

All of the experimental protocols involving animals were approved by the Children’s Hospital of Philadelphia Institutional Animal Care and Use Committee. B6 mice were bred in house or purchased from The Jackson Laboratory. β2 microglobulin^-/-^ mice were purchased from the Jackson Laboratory. Mice were infected with 3 × 10^3^ plaque forming units of ECTV by injection in a 30 μL volume of PBS through an insulin syringe into the hind footpad. All mice used for this study were females from 6-14 weeks of age, age matched within each study.

### Cells

NK cells used in *in vitro* assays were purified from spleens of B6 mice infected for 5-6 days with ΔC15 ECTV, using the EasySep mouse NK isolation kit (Stemcell, 19855). These cells were used directly in assays, in RPMI supplemented with 5% FBS, penicillin, streptomycin, 2mM L-glutamine and 50 μM 2-mercaptoethanol. The primary B6 skin fibroblast cell line was derived in our laboratory and has been described previously (Sinnathamby et al., 2004). These cells and TK-143b osteosarcoma cells were maintained in DMEM supplemented with 5% FBS, penicillin, streptomycin and 2mM L-glutamine. BSC-1 cells were maintained in DMEM supplemented with 10% FBS, penicillin, streptomycin and 2mM L-glutamine.

### Viruses

All viruses used in this work have been previously described: ECTV expressing eGFP (Moscow strain background) was a kind gift of Dr. Luis Sigal (Fang et al., 2008). eGFP expressing ΔC15 ECTV (Moscow strain background) was generated as previously described (Forsyth et al., 2020); the C15 revertant ECTV that was constructed was demonstrated to behave similarly to WT ECTV and was not used in this work. All viruses were grown in house in TK-143b osteosarcoma cells at high MOI for 3 days and virus was harvested from cells by cycles of freeze-thawing and sonication. Virus was then purified from the cell lysate by ultracentrifugation (20,000 rpm, 1 hour) through a 36% sucrose cushion and resuspended in 10 mM Tris at pH 9.0. To ensure titers were comparable, all viruses were titered simultaneously by traditional plaque assay on BSC-1 cells under an overlay of 1% methylcellulose in DMEM media supplemented with 5% FBS, penicillin, streptomycin and L-glutamine for 5 days.

### Viral titering in tissue

Mice were infected in the hind footpad for 1-3 days with 3000 pfu of eGFP WT or ΔC15 ECTV for 1-3 days. The whole draining popliteal LN was collected, weighed and homogenized. LNs were homogenized by manual disruption between two frosted glass slides and homogenates were then sonicated to release virions and cellular debris was pelleted by spinning at 400 xG for 10 seconds. The homogenate supernatant was titered by serial dilution on BSC-1 cells utilizing either a traditional plaque assay, as described above, or a focus forming assay as first described by Forsyth (Forsyth et al., 2020): 2.5×10^4^ BSC-1 cells were plated per well in a flat 96-well plate. The next day, serially diluted homogenates were added to monolayers in technical triplicate. Virus was allowed to adhere for 1 hour and then an overlay of 1.25% avicel in complete DMEM was added and infection was left to proceed for 18 hours. Then virus was removed, cell monolayers were fixed for 1 hour using 4% paraformaldehyde in PBS, permeabilized for 7 minutes using 0.5% Triton X-100 in PBS and then blocked for 1 hour with 5% BSA in Tris-buffered saline with 0.2% Tween 20. Virus infected cells were detected by staining with a primary rabbit anti-VACV antibody (1:1000 in blocking buffer for 1 hour, Thermo Fisher PA1-7258) and a secondary horseradish peroxidase-conjugated goat anti-rabbit antibody (1:1000 in blocking buffer for 1 hour, Cell Signaling Technology 7074S) with the peroxidase substrate KPL TrueBlue (30 minutes, SeraCare 5510-0030). Virus-infected foci representing one Focus Forming Unit (FFU) were imaged and counted spots using a CTL ImmunoSpot S6 Universal Analyzer with ImmunoSpot software (ImmunoSpot). When results were pooled from separate animal experiments, all samples were re-titered together on the same day with the same settings to account for any variation in the titering assay.

### In vivo NK cell depletion

B6 mice were depleted of NK cells by i.p. injection of 200 μg of anti-NK1.1 mAb (PK136, BioXCell BE0036) in 100 μL of PBS, compared to injection of IgG2a isotype control mAb (C1.18.4, BioXCell BE0085), 24 hours prior to infection. Depletion was confirmed by flow cytometric analysis of splenocytes for viral titering experiments and visually for immunofluorescence imaging experiments.

### Confocal microscopy of frozen LN sections

Popliteal LNs were harvested 3 days after infection with eGFP WT or ΔC15 ECTV, following isotype control or NK1.1 depletion. LNs were fixed overnight in periodate-lysine-paraformaldehyde buffer, equilibrated in 30% sucrose in PBS overnight and then embedded in Optimal Cutting Temperature medium (Sakura 4583) and frozen in liquid nitrogen cooled isopentane, as previously described (Reynoso et al., 2019). Sixteen micron sections were cut and blocked in .01% Triton X 100 with 2% of FBS and donkey serum including TrueStain Monocyte Blocker (BioLegend 426101) for 1h and then stained overnight with the following directly conjugated mAbs: GFP AF488 (clone FM264G, BioLegend 338007), NK1.1 AF647 (clone PK136, BioLegend 108719), Lyve-1 eFluor 450 (clone ALY7, Invitrogen 48-0443-80), ER-TR7 AF594 (Santa Cruz Biotechnology sc-73355), B220 AF700 (clone RA3-6B2, BioLegend 103232). Slides were washed in .1% Tween20 in PBS, mounted using ProLong Glass Antifade Mountant (Invitrogen P36984) and imaged using a Leica TCS SP8 WLL confocal microscope. Images were acquired using identical laser and HyD detector settings and scans were taken of an entire popliteal LN section using a 40x 1.30 NA objective, with a z step of 1.5 μm and a total z size of 9.0 μm; individual fields were then merged into a single image.

### Image Analysis

Raw data were processed using a gaussian filter to remove background noise, then processed into 3D maximum intensity projection. Spots were created for the NK cell (NK1.1+) channel using the automated Imaris function “spots” with manual correction as needed. Lymph node surface was created using the Lyve-1 channel. Surface for the virus infected cells was created based on the eGFP channel. The number of NK cells was calculated based on the number of spots generated. To calculate distance to the nearest neighbors, Imaris XT plugin “nearest neighbor” was applied to the NK spots and the average value was extracted from each image. The number of NK cells in contact with virus infected cells was calculated using the Imaris XT plugin “distance spots to surface” with a filtered distance of 5 μm or less between NK cells and virus-infected cell surface. Statistics were exported from Imaris. NK cell numbers were normalized to lymph node area. Values were then plotted in Prism (GraphPad).

### Flow cytometric analysis of infected tissue

Popliteal LNs were harvested 3 days after infection with eGFP WT or ΔC15 ECTV and incubated at 37°C in media containing monensin (Invitrogen 00-4505-51) and PE labeled CD107a antibody (1:100, clone EDB4, BioLegend 121612) for one hour. LNs were then dissociated using frosted glass slides, filtered through a 40 μm strainer, and entire LN cell populations were counted and stained for flow cytometric analysis. Cells were first stained in Live/Dead Blue (1:500, Invitrogen L23105), treated for Fc-receptor blockade with anti-CD16/32 (100 μg/mL, BioXCell BE0307) and then stained for surface antibodies at .1 μg/mL unless indicated: NK1.1 PerCP-Cy5.5 (clone PK136, BioLegend 10727), F4/80 AF594 (clone BM8, BioLegend 123140), CD3e APC-Cy7 (clone 145-2C11, BD 557596). Cells were then fixed and permeabilized for intracellular staining with the BD Cytofix/Cytoperm kit (BD 554714) and then stained for intracellular antibodies at .1 μg/mL unless indicated: GzB BV510 (1:10, clone GB11, BD 563388), Ki-67 APC (clone SolA5, Thermo Fisher 17-5698-82), IFNγ AF700 (clone XMG1.2, BD 557998). Flow cytometry data were acquired on the CytoFLEX LX instrument using CytExpert software (Beckman Coulter) and analyses were conducted using FlowJo software (FlowJo LLC). NK cells were identified as CD3^-^NK1.1^+^ live singlets. Median fluorescence intensity (MFI) was analyzed for the positive staining populations.

### RNA isolation and RT-PCR

Total RNA was extracted from LNs by bead homogenization and purified using the Qiagen RNeasy Plus Mini Kit (Qiagen 74104) and cDNA was prepared from 750 ng of RNA using the High Capacity cDNA Reverse Transcription Kit (Applied Biosystems 4368814). Quantitative PCR was performed using the PowerUp SYBR Green Master Mix (Applied Biosystems A25742) and measured using a StepOnePlus Real Time PCR Machine with StepOnePlus software (Applied Biosystems). Expression was quantified relative the housekeeping gene *Gapdh* normalized to the mean of the mock-infected group, using the ΔΔC_T_ method. Fold change values, 2^-ΔΔCt^, were pooled from 3 independent experiments. The forward and reverse primer pairs used are listed here: Quantitative PCR primers

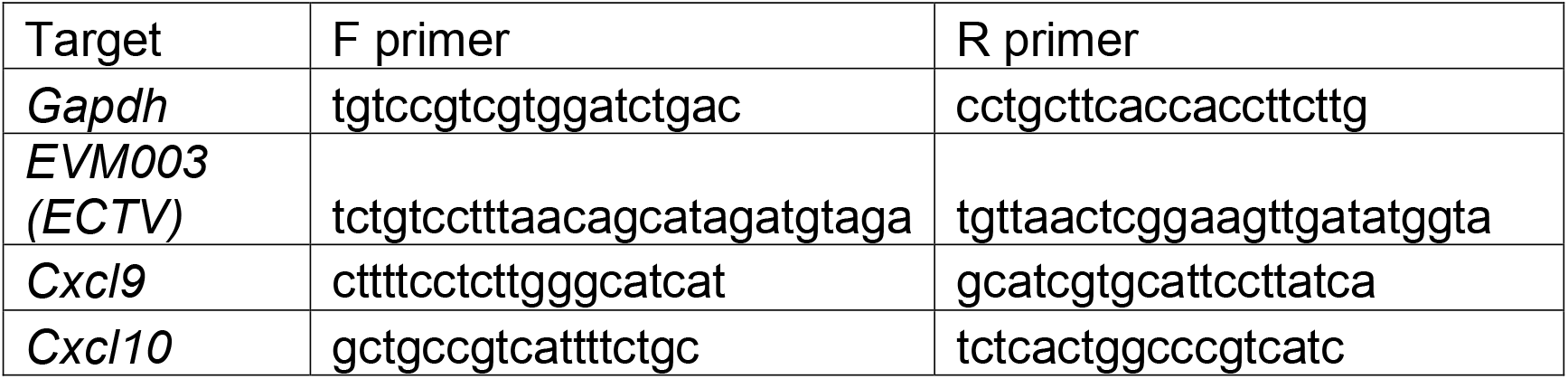
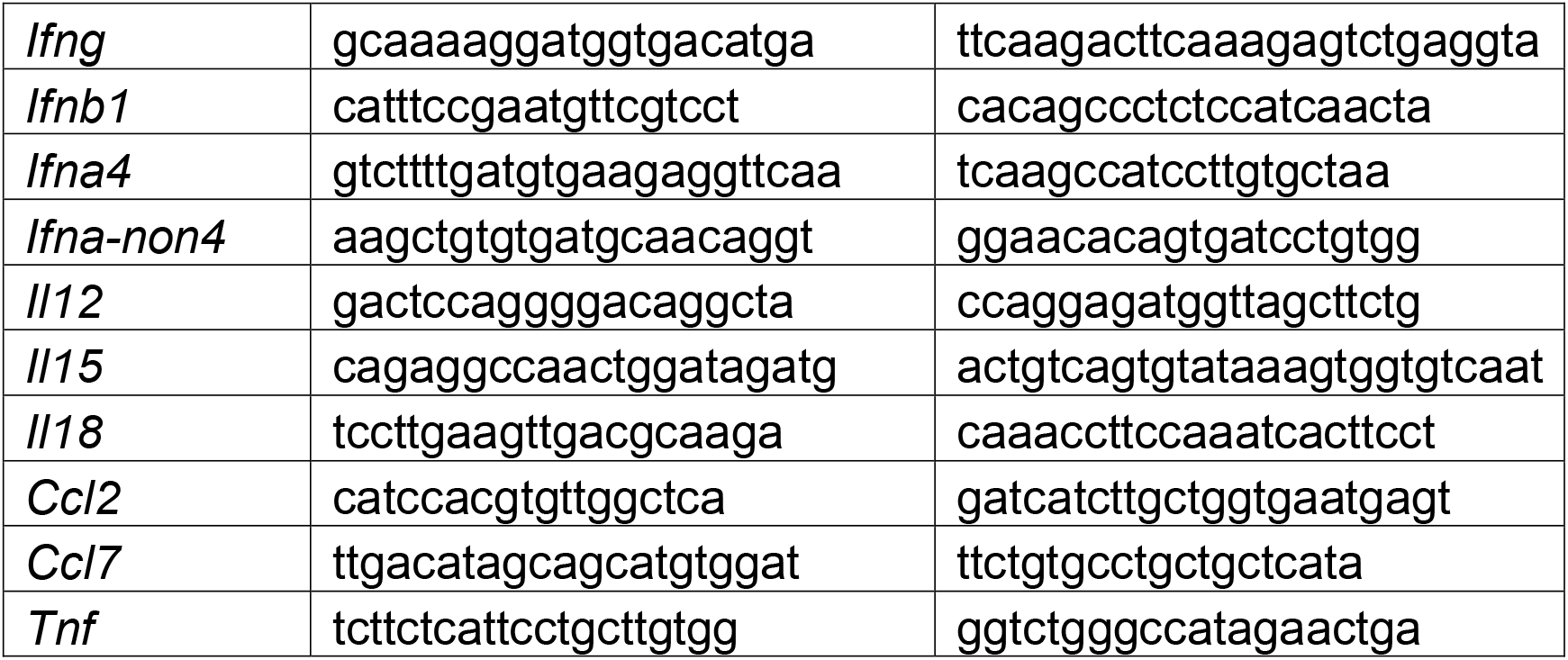

### In vitro NK degranulation assay

Primary B6 skin fibroblast cells were mock infected with PBS or were infected with an MOI of 3 of eGFP WT or ΔC15 ECTV for 1 hour in minimal volume and then infection was allowed to proceed for a total of 6 hours. At this time, these target cells were plated at the indicated effector:target ratios with NK cells that were isolated as described above. These cocultures proceeded for 5 hours in the presence of monensin (Invitrogen 00-4505-51) and PE labeled CD107a antibody (1:100, clone EDB4, BioLegend 121612), and in the indicated experiments with 20 μg/mL of blocking anti-NKG2D antibody (R&D systems MAB1547). Cells were then washed, stained for Live/Dead Blue (1:100 Invitrogen L23105) and extracellular antigens (1:200 CD3 BV650, clone 145-2c11, BD 564378; 1:200 NK1.1 PerCP-Cy5.5, clone PK136, BioLegend 10727) for 30 mins in PBS, fixed and permeabilized using the BD Cytofix/Cytoperm kit (BD 554714) and then stained intracellularly (1:200 IFNγ AF700, clone XMB1.2, BD 557998) for one hour. Data were acquired on the CytoFLEX LX instrument using CytExpert software (Beckman Coulter) and analyses were conducted using FlowJo software (FlowJo LLC). NK cells were identified as CD3^-^NK1.1^+^ live singlets.

### Statistical analyses

Statistics analyses were performed in Prism 9. Wherever possible, all data from three experimental replicates is pooled for display and analysis. Data were tested for normality, using the D’Agostino & Pearson test, prior to selecting parametric analyses. In instances in which an outlier test revealed that non-normality was driven by a single data point, the data was considered normal and parametric tests were selected. Where possible, lognormal data was log transformed to enable use parametric tests. Any manipulation of the data, such as log transformation, and the statistical test used are indicated in the figure legend and p values are shown for transparency. Individual biological or technical repeats are shown as points for transparency and all bars correspond to means and error bars represent SEM.

## Non-Standard Abbreviations

CPXV: cowpox virus
dpi: days post infection
ECTV: ectromelia virus
eGFP: enhanced green fluorescent protein
FFU: focus forming units
GzB: granzyme B
LN: lymph node
MOI: multiplicity of infection
MPXV: monkeypox virus
MFI: median fluorescence intensity
NK: natural killer
OPXV: orthopoxvirus
ORF: open reading frame
PFU: plaque forming units
TI-IFN: type 1 interferons
VACV: vaccinia virus
VARV: variola virus
ΔC15: C15-deficient eGFP expressing ectromelia virus

## Online Supplemental Material

Fig. S1 shows the concordance of our focus forming assay with a traditional plaque assay and demonstrates the extent of our NK cell depletion. Fig. S2 shows the full set of images used in Figs. 1 C and D to demonstrate the spectrum of viral replication and spread. Fig. S3 shows that C15 does not rely upon NKG2D to inhibit NK cell degranulation *in vitro* but that C15 does require β2 microglobulin to facilitate early replication in the LN.

## Acknowledgements

We thank the CHOP Department of Veterinary Resources and Flow Cytometry Core, the Penn Vet Comparative Pathology Core and the Penn Medicine CDB Microscopy Core. We also thank Drs. David Christian and Christopher Hunter for assistance with microscopy and Drs. Leilani Chirino and Taku Kambayashi for assistance with NK assays.

## Author Contributions

Conceptualization: Elise M. Peauroi, Stephen D. Carro, Heather D. Hickman, Laurence

C. Eisenlohr.

Formal analysis: Elise M. Peauroi, Luxin Pei.

Funding acquisition and project administration: Elise M. Peauroi, Laurence C. Eisenlohr.

Investigation: Elise M. Peauroi.

Methodology: Elise M. Peauroi, Glennys V. Reynoso, Luxin Pei.

Supervision: Laurence C. Eisenlohr, Heather D. Hickman.

Validation: Elise M. Peauroi, Luxin Pei.

Visualization: Elise M. Peauroi, Luxin Pei.

Writing – original draft: Elise M. Peauroi.

Writing – review & editing: Elise, M. Peauroi, Stephen D. Carro, Luxin Pei, Glennys V.

Reynoso, Heather D. Hickman, Laurence C. Eisenlohr.

The authors declare no competing financial interests.

This work was supported by National Institutes of Health grants R21AI160063 (to Laurence C. Eisenlohr), T32AI070077 (to Michael L. Atchison) & F30AI149864 (to Elise M. Peauroi).

---

**Figure S1.**
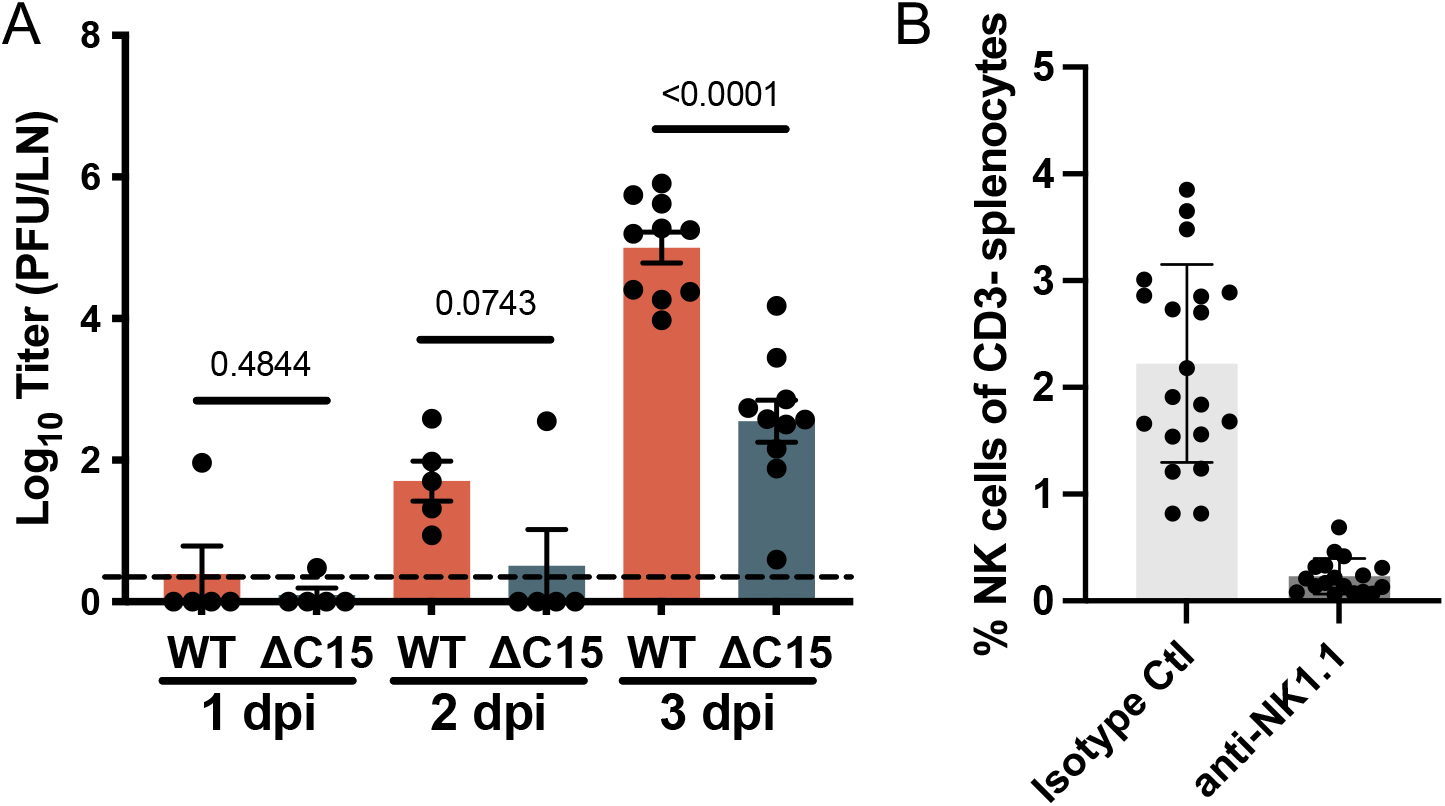
Success of focus forming assay and NK depletion. **(A)** The same tissues as in Fig. 1 A were analyzed by traditional plaque forming assay to confirm that conclusions are similar by FFA. Graph displays log_10_ transformed titers, as plaque forming units (PFU) per LN. **(B)** Depletion confirmation from the mice in Fig. 1 B. At sacrifice, the spleen was collected, processed and splenocytes were analyzed by flow cytometry to confirm NK cell depletion. NK cells were gated as CD3-NK49b+NKG2D+ live cells. NK cells are presented as percent of CD3-splenocytes. Bars = mean, Error bars = SEM. T tests were performed; p values are shown.

**Figure S2.**
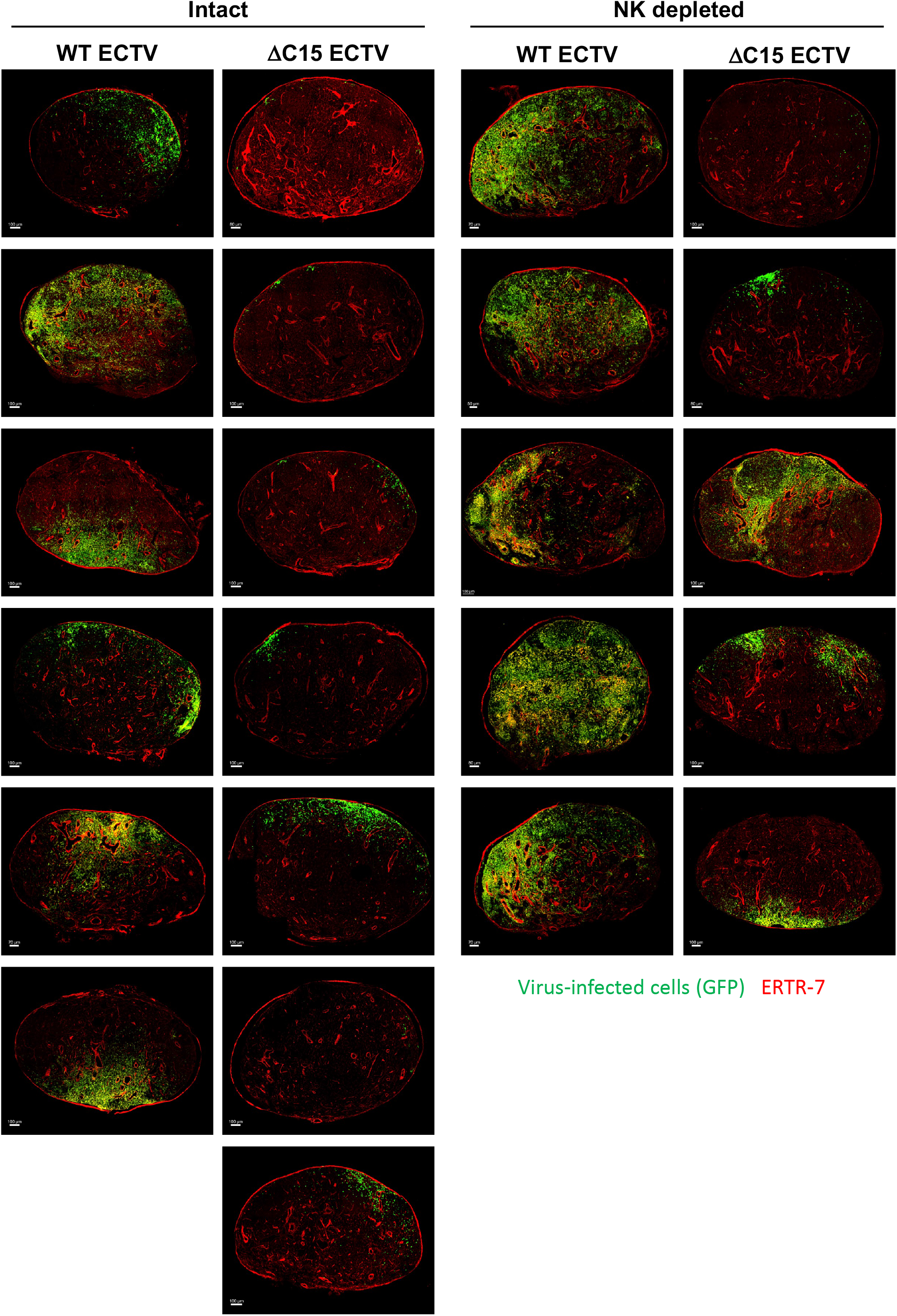
Full data set of images demonstrating the variation in viral dissemination in the LN. Draining popliteal LNs were harvested 3 dpi with WT or ΔC15 and analyzed for the indicated markers by immunofluorescence imaging. The demonstrated images are maximum intensity projections from each biological replicate. Scale bars are labeled. Images are pooled from 3 independent experiments.

**Figure S3:**
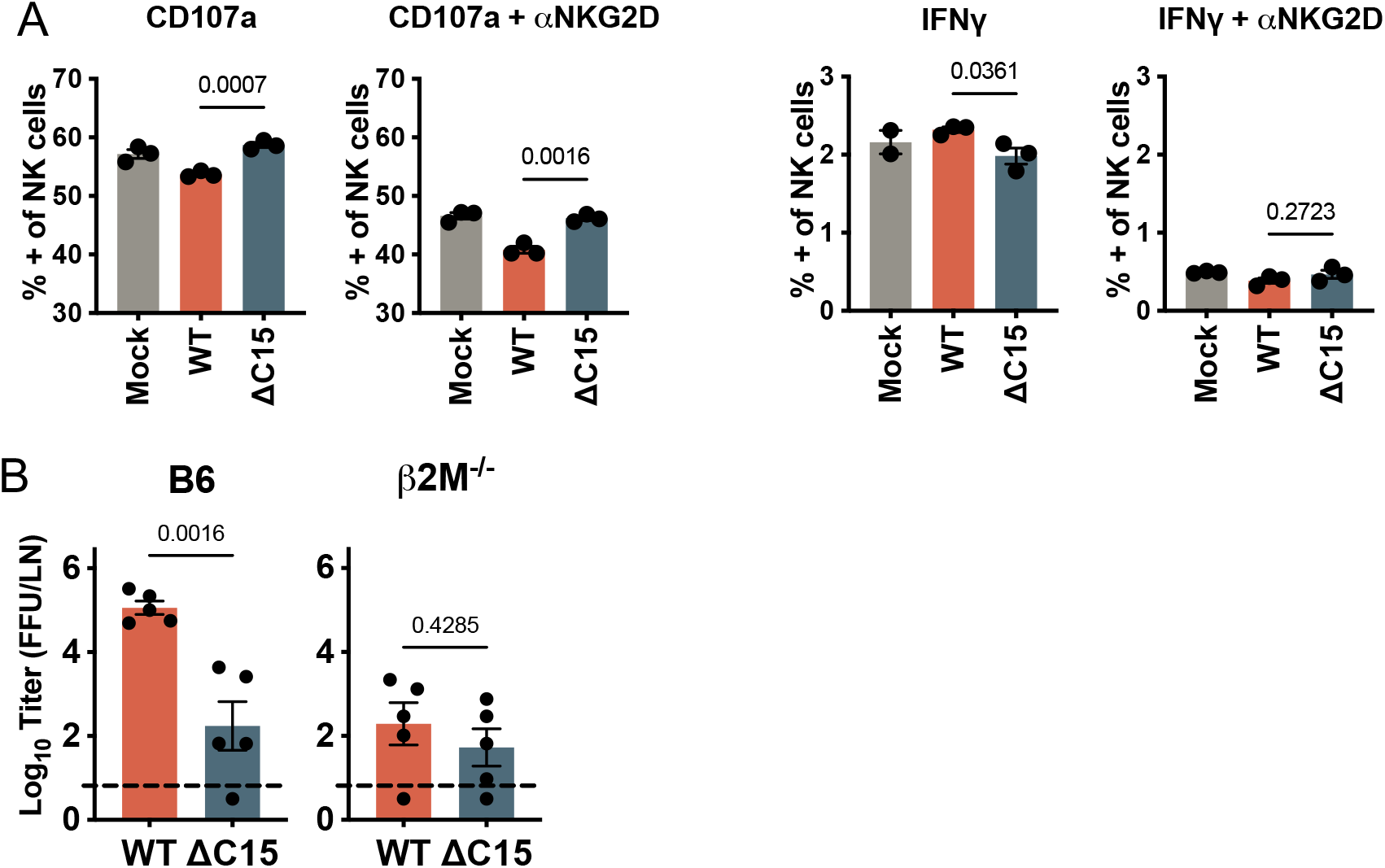
C15 appears to rely upon β2 microglobulin and not NKG2D. **(A)** The *in vitro* NK cell degranulation experiment in Fig. 4A was conducted in the presence or absence of anti-NKG2D blocking antibody, and again degranulation (surface CD107a) and IFNγ production (intracellular) were analyzed. Data points are technical replicates, bars represent means and error bars are SEM. **(B)** B6 and β2 microglobulin deficient mice (β2M^-/-^, n=5, single experiment) were infected with 3000 pfu of WT or ΔC15 ECTV and sacrificed 3 dpi. The draining popliteal LNs were harvested, homogenized and viral titers were determined by focus forming assay on BSC-1 cells. Graphs display log_10_ transformed titers as FFU/organ. Bars = mean, Error bars = SEM. T tests were performed: p values are shown. Both A and B represent single experiments.

